# Genomic Epidemiology and Microevolution of the Zoonotic Pathogen *Corynebacterium ulcerans*

**DOI:** 10.1101/2024.08.22.609154

**Authors:** Chiara Crestani, Virginie Passet, Martin Rethoret-Pasty, Alexis Criscuolo, Nora Zidane, Sylvie Brémont, Edgar Badell, Sylvain Brisse

**Affiliations:** Institut Pasteur, Université Paris Cité, Biodiversity and Epidemiology of Bacterial Pathogens, Paris, France; Institut Pasteur, National Reference Center for Corynebacteria of the diphtheriae complex, Paris, France; Institut Pasteur, Biological Resource Center of Institut Pasteur, Paris, France

**Keywords:** *Corynebacterium ulcerans*, genomic epidemiology, diphtheria toxin, transmission, cgMLST

## Abstract

*Corynebacterium ulcerans* is an emerging zoonotic pathogen that belongs to the *Corynebacterium diphtheriae* (Cd) species complex (CdSC), and that causes diphtheria-like infections in humans. Our understanding of the transmission, phylogeography and evolution of *C. ulcerans* remains limited, in part due to the lack of a standardized genomic epidemiology toolkit.

The aim of this work was to develop a core genome multi-locus sequence typing (cgMLST) scheme for high-resolution genotyping and classification of *C. ulcerans* strains, and to explore transmission, spatial spread and genomic evolution among 582 *C. ulcerans* isolates from sporadic clinical cases and reported case clusters.

The cgMLST scheme combines 1,628 loci with highly reproducible allele calls and shows high strain subtyping resolution. We demonstrate its utility for capturing population structure by defining sublineages (SL, maximum 940 allele differences) and clonal groups (CG, 194 allele differences, AD) and for epidemiological surveillance by defining genetic clusters, i.e., previously undetected chains of transmission (25 AD). Genetic clusters correspond to cryptic and case clusters that were associated with specific geographical regions within France. Major *C. ulcerans* sublineages (SL325, SL331, SL339) and clonal groups (CG325, CG331, CG583) showed strong associations with diphtheria toxin variants and tox-carrying prophages or other genetic elements. The evolutionary dynamics of *tox* gene presence or absence varied sharply among clonal groups. The cgMLST scheme is publicly available (https://bigsdb.pasteur.fr/diphtheria) and provides a common framework for investigating the ecology, evolution and variations in virulence among *C. ulcerans* strains. The implementation of a standardized high-resolution genotyping method will also facilitate the tracing of *C. ulcerans* transmission and spread across hosts and from local to global spatial scales.

## Introduction

*Corynebacterium ulcerans* is a member of the *Corynebacterium diphtheriae* Species Complex (CdSC) and a cause of diphtheria-like respiratory infections in humans as well as cutaneous infections (Moore et al., 2015; Othieno et al., 2019; Otshudiema et al., 2021)*. C. ulcerans* isolates can carry the *tox* gene coding for diphtheria toxin, a prominent virulence factor responsible for systemic clinical signs that worsens the prognosis (Hacker et al., 2016; Mueller, 1941). Among *C. ulcerans* isolates collected at the French national reference centre (NRC) for diphtheria, a high proportion are toxin-positive isolates (56%; data for 2023 from the French NRC). The toxin gene can be carried by prophages, as in *C. diphtheriae,* and additionally by a pathogenicity island (PAI) described so far only in *C. ulcerans* (Dangel et al., 2019; Freeman, 1951; Parveen et al., 2019).

In the last decade, human *C. ulcerans* infections have been on the rise in several countries (A. A. Dias et al., 2011; Hacker et al., 2016; Moore et al., 2015; Wagner et al., 2010), and most cases can be traced to contact with diseased or healthy pets (e.g., dog and cats) (Corti et al., 2012; Hoefer et al., 2022; Meinel et al., 2015; Saeki et al., 2015; Simpson-Louredo et al., 2014). *C. ulcerans* has been reported in multiple animal species, including domestic (e.g., cattle, pigs, goats) (Boschert et al., 2014; Higgs TM et al., 1967; Hommez et al., 1999; Morris et al., 2005; Museux et al., 2023) and wild animals (e.g., hedgehogs, wild boars, roe deer) (Berger et al., 2019; Eisenberg et al., 2014; Martel et al., 2021).

Little is known about the transmission dynamics of *C. ulcerans* between these species, especially between wildlife and domestic animals, limiting our ability to control infections in animals and zoonotic transmission. Besides, the phylogeography of *C. ulcerans,* i.e., how genotypes disseminate geographically, is unknown. Furthermore, the small-scale evolution of *C. ulcerans*, for example how frequently sublineages may gain or lose the *tox* gene, remains unexplored. This is in part due to the lack of inclusion of veterinary *C. ulcerans* isolates in most national surveillance programmes, but also to the lack of a standardized genotyping system that would enable to define and track *C. ulcerans* strains. Currently available genotyping methods for *C. ulcerans* include seven-gene multilocus sequence typing (MLST) (Maiden et al., 1998). However, this classical approach suffers from severely limited resolution and hence, an insufficient discriminatory power to detect case clusters or chains of transmission.

The advancements of whole genome sequencing (WGS) technologies and analytical methods have allowed for the expansion of the MLST method to the wider core genome (cgMLST), combining a high resolution with the well-established reproducibility and portability of MLST approaches. Species-specific cgMLST schemes have been developed for several bacterial pathogens, including *C. diphtheriae* and its closely related species *C. rouxii* and *C. belfantii* (Guglielmini et al., 2018), allowing for successful detection of cryptic transmission events and epidemiological clusters in this bacterial group. However, no cgMLST method is currently publicly available for *C. ulcerans*.

The aims of this study were: i) to provide a publicly available standard method for accurate and high-resolution *C. ulcerans* strain genotyping, and ii) to establish a baseline framework for investigating the ecology, evolution, transmission dynamics and variations in virulence across the population diversity of *C. ulcerans*. We developed and benchmarked a cgMLST scheme for *C. ulcerans*, showing it represents a highly accurate genotyping approach, and we investigate the population structure of *C. ulcerans* across continents, defining sublineages (SL) and clonal groups (CG), as well as genetic clusters that may reflect chains of transmission. The revealed population structure was then used to analyse the phylogenetic distribution and the evolutionary dynamics of the main virulence genes of *C. ulcerans,* including the diphtheria toxin gene, *tox*, and its associated mobile genetic elements.

## Materials and methods

### Bacterial isolates and case clusters included in the study

Three datasets were used for this study: i) an original dataset comprising 347 *C. ulcerans* genomes available in June 2022 from the French NRC and the Collection of Institut Pasteur (n=317), and from NCBI (n=30^1^); ii) an expanded dataset of 434 genomes, comprising additional *C. ulcerans* (n=51) and *C. ramonii* (n=36) isolates from private and public repositories; iii) and a validation dataset of 582 *C. ulcerans* genomes, which comprised additional genomes (n=184) sequenced at the French NRC while this work was in progress and publicly available *C. ulcerans* genomes from the *Corynebacterium diphtheriae* species complex BIGSdb database (Jolley & Maiden, 2010), including case clusters.

The first two datasets were used to build the core genome multi-locus sequence typing (cgMLST) scheme for *C. ulcerans*, that would be also applicable to *C. ramonii*, whereas the third dataset was used for validation of the cgMLST scheme, for sublineage (SL) and clonal group (CG) definition, as well as for exploration of case clusters and of clusters with cryptic transmission.

The latter dataset comprised 20 confirmed clusters for which epidemiological data was available (Table *1*). Six of these originated from France, one from Spain, five from Germany, one from the UK and seven from Belgium. The latter comprised four case clusters for which only one genome sequence was available.

**Table 1.**
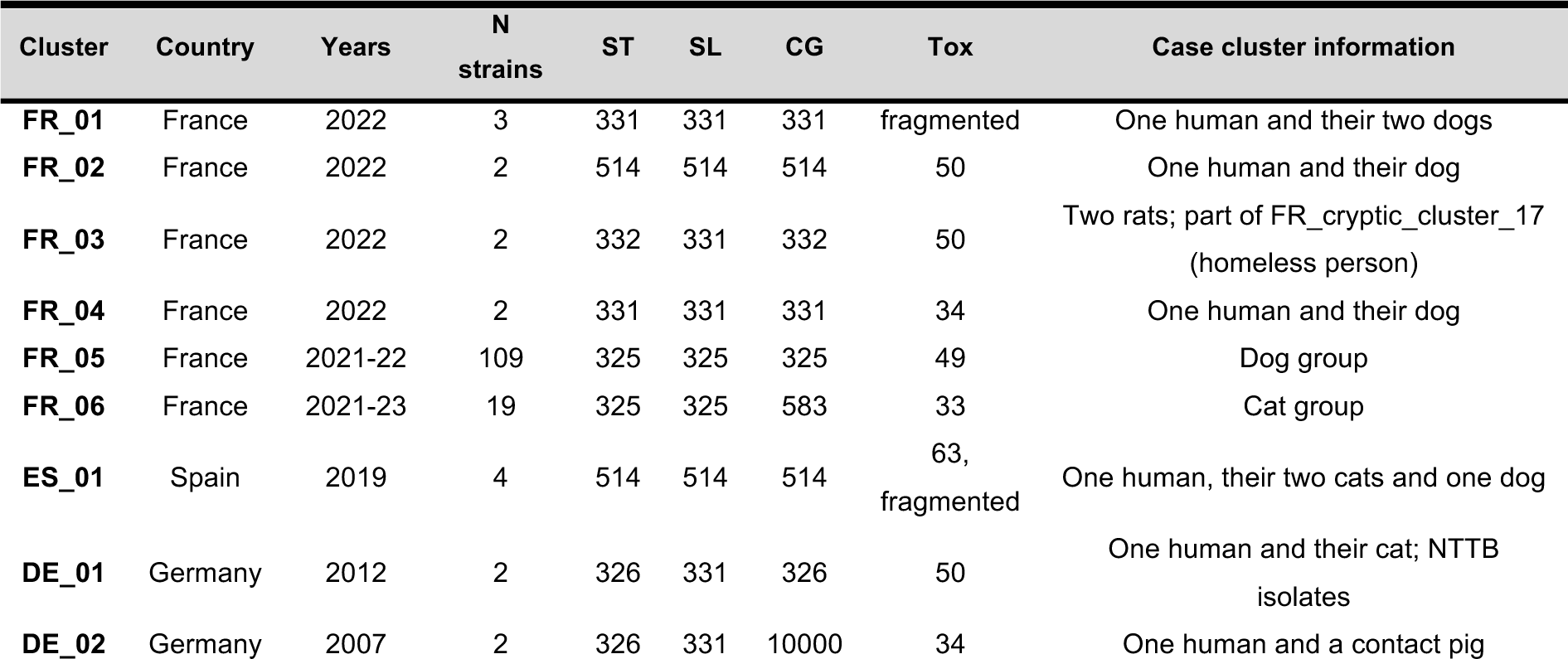
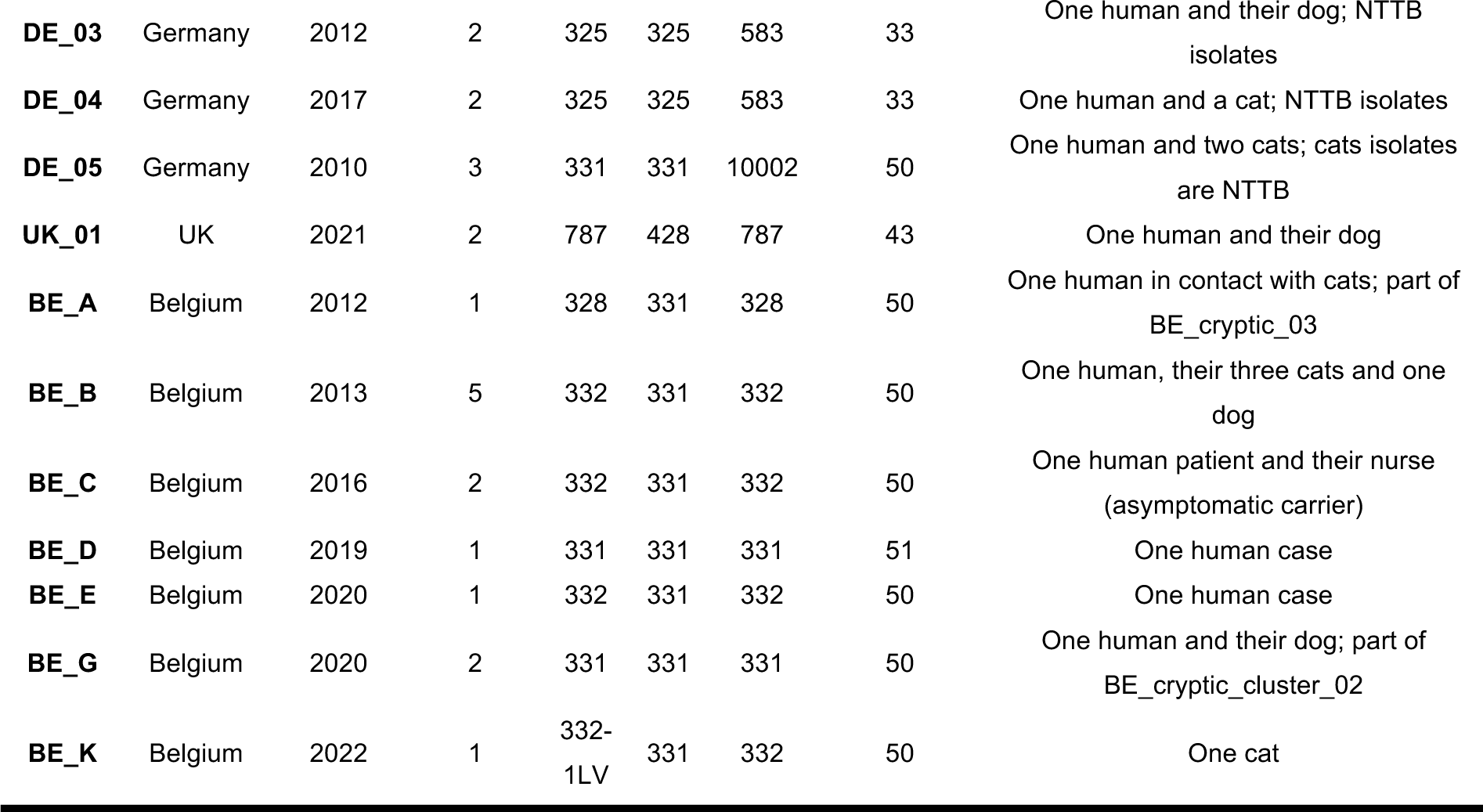
*Corynebacterium ulcerans* case clusters included in this study.

### Isolate growth and DNA extraction

Isolates were grown in Tryptic Soy Agar (TSA) for 24-48 hours at 35–37 °C and then resuspended in sterile saline (0.9% NaCl) for DNA extraction (optical density, OD=1), which was performed using the DNeasy Blood & Tissue Kit (QIAGEN, Hilden, Germany). A lysis step was added to the extraction protocol described by the manufacturer (Badell et al., 2019).

### Genomic sequencing and assembly

Whole Genome Sequencing (WGS) was performed from Nextera XT libraries on an Illumina NextSeq-500 apparatus (Illumina, San Diego, USA) with a 2 x 150 nt paired-end protocol.

Paired-read sequence data were assembled with the pipeline fq2dna v21.06 (https://gitlab.pasteur.fr/GIPhy/fq2dna), and assembly quality was checked with QUAST v5.0.2 (Gurevich et al., 2013). Only genomes with a GC% within the range of 53.33% ± 0.51% (corresponding to twice the standard deviation), and with a genome length within the range of 2,533,898 bp ± 180,868 bp (twice the standard deviation) were used for this study. Species identity based on Mash distances (Ondov et al., 2016), sequence types (ST) and the presence of the *tox* gene were determined with the software tool diphtOscan (Hennart et al., 2023).

### Definition of the cgMLST scheme

From a starting dataset of 347 genomes, we selected 239 as being representative of the diversity of *C. ulcerans*. These included representatives of all Sequence Types (ST) (n=38) that had ≥21 Single Nucleotide Polymorphisms (SNPs) in the core genes, except for FRC1398, which was also included, and which carries a toxin allele different from that of all other genomes in the same case cluster.

To build the cgMLST scheme, a dedicated tool for creation and validation of cgMLST schemes, chewBBACA v2.8.5 (Silva et al., 2018), was selected. We used the following functions available within the chewBBACA suite: i) Whole genome Multi-Locus Sequence Typing (wgMLST) with default settings (BLAST Score Ratio of 60%, CDS size variation threshold of 20%); ii) Allele call using the wgMLST scheme (paralogs are eliminated automatically at this step); iii) Evaluation of wgMLST call quality per genome; iv) Definition of the cgMLST scheme (95% threshold). The following user-defined options were implemented: a) A custom Prodigal training file (Culcerans_0102, the largest genome size and *tox* gene positive assembly available) was used to better identify CDS; b) Removal or repeated loci; c) Removal of genomes with low quality scores. The pipeline resulted in a cgMLST scheme of 1,941 loci, from which loci of the seven-gene MLST scheme and of the ribosomal MLST scheme were removed (total hits n=53), for a final count of 1,888 cgMLST loci. This set of 1,888 loci was compared to the loci of the existing cgMLST scheme for *C. diphtheriae* (n=1,305 loci) with BLASTn (word size=10, e-value≤0.0001), using alleles from the type strains of the two species (NCTC7910^T^ for *C. ulcerans* and NCTC11397^T^ for *C. diphtheriae*). With thresholds set at >69% for identity and ≥70% for query coverage (chosen based on the distributions of these two variables, Figure S1, Figure S2), and allowing a 5% allele size variation, 519 loci were found to be in common between the two schemes. These make up 40% and 30% of the *C. diphtheriae* and *C. ulcerans* cgMLST schemes, respectively.

The expanded dataset (n=434), which included 36 isolates of lineage 2 recently described as *C. ramonii*, was first tagged for the 1,888 cgMLST loci (with the autotag.pl script and the following parameters: -w 30, -f), then scanned for new alleles (with the scannew.pl script and the following parameters: -w 30, - f, -c, 90% query coverage and 90% identity), and finally re-tagged to attribute novel alleles to the genomes. A total of 721,064 out of 819,392 maximum possible CDS (88%) were identified, with n=251 loci showing ≥10% of missing data. These were therefore removed from the scheme, leaving a set of 1,637 remaining loci.

Finally, to filter out loci with non-reproducible allele calls at low assembly coverage depths, randomly selected read pairs simulating various coverage depths (10X, 20X, 30X, 50X, 80X, 100X) were drawn from four genomes (FRC0027, FRC0058, FRC0687, FRC0804; in triplicate for each genome at each coverage). All alleles were called with no mismatches at 80X and 100X, whereas 9 unique loci showed non-reproducible allele calls at 30X and 50X (Figure S3). These were eliminated from the scheme, leaving a final set of 1,628 loci, which are likely to be present in most isolates of *C. ulcerans* and *C. ramoniii*, single copy with no paralogs, and called with precision even at moderate sequence read depths.

### Implementation of the cgMLST scheme for *C. ulcerans* in the *Corynebacterium diphtheriae species* complex BIGSdb database

The reference alleles for the 1,628 loci cgMLST scheme were extracted from the NCBI reference genome for *C. ulcerans*, strain 809, and they were reordered based on the position of detection on the reference genome (5’ end of each CDS). Loci shared with the cgMLST of *C. diphtheriae* were not renamed (n=497), whereas loci that were unique to the scheme of *C. ulcerans* (n=1,131) were renamed based on their position in the reference genome, incrementally (e.g., ULC_0001, ULC0002 etc.). Type alleles for *C. ulcerans,* which are used at the scannew step to search for novel alleles, were registered for those loci (n=497) in common with the *C. diphtheriae* scheme. These were then incorporated in the *Corynebacterium diphtheriae* species complex BIGSdb database (https://bigsdb.pasteur.fr/diphtheria) as a dedicated scheme (cgMLST_ulcerans).

All genomes in the expanded dataset (n=434) were tagged on BIGSdb for 1,628 loci (autotag.pl script, see parameters above), and only those sequenced with Illumina technology (n=420) were used to define new alleles with the scannew.pl script. Then, the whole set of genomes was tagged again, and cgMLST profiles were defined with the define_profiles.pl script, tolerating a threshold of 5% missing alleles (n≤81) to define cgSTs in the seqdef database. The same process was repeated to call alleles and define profiles in the additional isolates (n=235) of the third dataset.

### Cryptic cluster detection

Cryptic clusters were defined as single-linkage groups of genomes based on a threshold of ≤25 cgMLST allelic mismatches, for which no prior information on epidemiological links was available.

### Phylogenetic and population structure analyses

A core gene alignment obtained with panaroo (default settings) was used for phylogenetic inference with IQ-TREE v2.0.6 (GTR+F+R8) (Minh et al., 2020; Nguyen et al., 2015). Four *Corynebacterium pseudotuberculosis* genomes from reference strains (ATCC19410, CCUG27541, NCTC4656, NCTC4681) were used as outgroup to root the tree. The tree was visualised with Microreact (Argimón et al., 2016). Figures were edited using Inkscape (available from: https://inkscape.org).

### Toxin gene, and *tox*-associated prophage and pathogenicity island

The diphtheria toxin gene was detected on available isolates by a qPCR assay (Badell et al., 2019), and the production of the toxin was evaluated with the modified Elek immunoprecipitation test (Engler et al., 1997).

The presence of the toxin gene and of other *C. ulcerans*-specific virulence genes (e.g., *pld, rbp*) was assessed in WGS data from the validation dataset with the software diphtOscan (Hennart et al., 2023). When a new toxin allele was found, the nucleotide sequence was further explored via the BIGSdb-Pasteur platform to ensure it corresponded to a CDS, and subsequently added to the toxin allele database.

All *tox*-positive genomes were scanned with BLAST+ (Camacho et al., 2009) for the presence of the toxin-encoding pathogenicity island (PAI, reference sequence available at GenBank under accession n. KP019622), and with PHASTER (Arndt et al., 2016) for the presence of toxin-carrying prophages. *Tox*-prophages sequences were then manually curated with Geneious Prime 2024.0. When *tox*-positive genomes were negative for the presence of either the PAI or prophages, manual exploration of these was carried out with SnapGene Viewer v7.1.0 (https://www.snapgene.com). Orphan toxins (i.e., present as a single gene contig) were detected consistently among CG325 genomes (see below); therefore, three genomes in this clonal group (FRC1215, FRC1277, FRC1308) were selected for sequencing using Oxford Nanopore MinION R9.4.1 flow cells with the Rapid Barcoding kit (SQK-RBK004), to clarify the location of the toxin (see results section). Basecalling was performed with Guppy v6.4.2 (https://nanoporetech.com/) using the SUP algorithm, and high-quality hybrid Illumina-Nanopore assemblies were obtained with Unicycler v0.4.8 (Wick et al., 2017). To reconstruct the *tox-*prophage from the remaining genomes of the CG325, raw reads were mapped to the reference prophage of FRC1215 with bwa-mem2 v2.2.1 (Li, 2013), and consensus sequences were obtained with samtools v1.18 (Danecek et al., 2021).

Multiple sequence alignments were performed for: i) all toxin alleles in the BIGSdb-database (n=63); ii) full prophage sequences (n=176 from *C. ulcerans* and n=232 from *C. diphtheriae*); and iii) for the PAI (n=189) using mafft v7.526 (Katoh & Standley, 2013) for the former, and Clustal-Omega v1.2.4 (Sievers et al., 2011) for the latter two, with default settings (Gap opening penalty = 10, Gap extension penalty = 0.20). Integrase genes were extracted from prophage sequences with BLAST+.

Approximately-maximum-likelihood phylogenetic trees for both the toxin alleles and the corresponding *tox-*prophages were constructed from nucleotide sequence alignments using FastTree v2.1.11 (Price et al., 2010) using the Jukes–Cantor model with default parameters. Phytools v2.1-1 (Revell, 2012) was used to obtain a co-phylogeny of the toxin alleles and the prophage sequences.

## Results

### Definition and evaluation of a cgMLST scheme for genotyping of *C. ulcerans* and *C. ramonii*

We defined a set of 1,628 protein-coding genes for genotyping of *C. ulcerans* and *C. ramonii* and combined them into a cgMLST scheme. Locus length varied from 204 to 9,099 bp, and all loci combined had a total length 1,682,922 bp (calculated for allele 1 of each gene), representing 69% of the genome size of the reference strain NCTC7910^T^.

The allele call rate for *C. ulcerans* was defined based on 235 genomes that were not included in the construction of the cgMLST scheme, including genomes from Belgium (n=34), Germany (n=47), Japan (n=5), Norway (n=1), Spain (n=4), the UK (n=3), unknown geographical origin (n=2), and newly-sequenced genomes from the French NRC (n=139). The allele call rate was found to be 99.7% ± 0.2%, with a mean number of called alleles of 1,623 per genome. For *C. ramonii*, as this species is rare, only four additional genome sequences that were not included in the dataset for cgMLST building were publicly available. Therefore, for scheme validation, we calculated the allele call rate for all *C. ramonii* isolates available (n=40, n=36 of which from the second dataset), excluding those that have been sequenced with technologies other than Illumina (n=5), showing an allele call rate of 96.5% ± 0.9% and an average of 1,571 called alleles per genome.

Based on the 582 genomes of the overall study dataset, each locus has between 2 to 91 unique alleles, with the number of alleles increasing with locus size (Figure S4). The longest locus was atypically long (9,099 bp), coding for a type I polyketide synthase, and had the highest number of alleles as expected from the positive correlation between locus size and allele number.

### Population structure of *C. ulcerans*

The phylogenetic structure of the *C. ulcerans* population showed a few deep branches and many clusters of highly similar genomes, which is indicative of clonal expansions (e.g., CG331 or CG339, see below) corresponding, in a few cases, to documented outbreaks (e.g., CG325) (Figure *1*).

**Figure 1.**
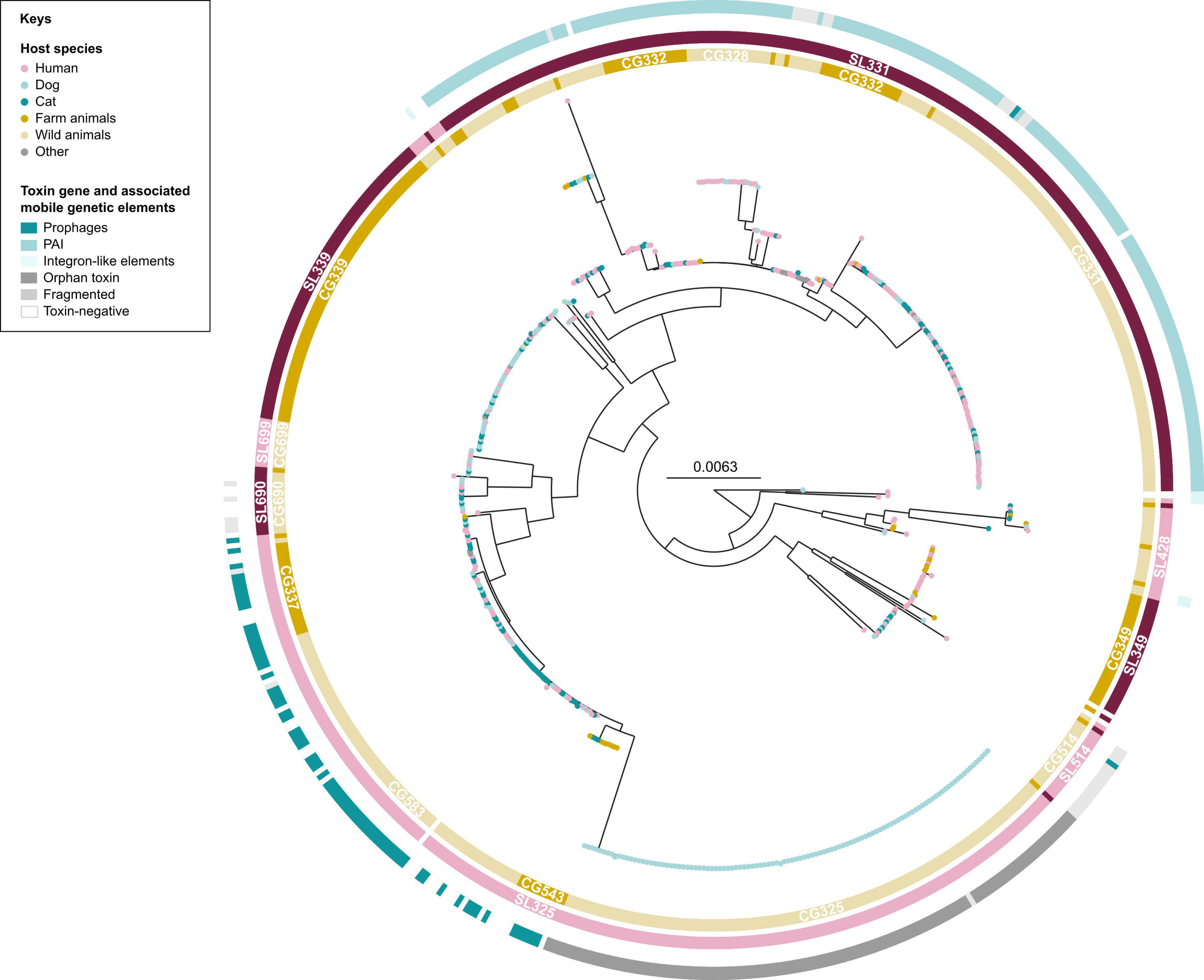
Maximum-likelihood phylogenetic tree of 582 *Corynebacterium ulcerans* from human and animal isolates. Coloured leaves indicate the host species (human, dog, cat) or host group of origin (farm animals, wild animals). External strips show the clonal groups (CG), with their respective number inherited from the seven-gene MLST nomenclature for the most important ones, and the sublineages (SL). For all *tox-*positive isolates, the associated mobile genetic element carrying the diphtheria is shown in the outermost strip, when its identification was possible (e.g., except for fragmented genomes). The tree was rooted on *Corynebacterium pseudotuberculosis* (collapsed).

MLST data from the 7-gene scheme showed 54 different STs, with four being very common: ST325 (n=179, 31%), ST331 (n=99, 17%), ST339 (n=62, 11%) and ST332 (n=51, 8.8%). These comprised 67.2% of isolates in the validation dataset (unless otherwise stated, we use the validation dataset for epidemiological inferences hereafter). STs were not always concordant with the phylogenetic structure of *C. ulcerans* (Microreact project), as ST326, ST331 and ST332 were polyphyletic; however, all other non-singleton STs were monophyletic. ST325 was monophyletic but highly heterogeneous, being subdivided into four different clonal groups (CG325, CG583, CG543, CG10001; see below).

To identify optimal thresholds of allelic differences among cgMLST profiles that would define genetic clusters most-fitting to the *C. ulcerans* population structure, we calculated the consistency and stability properties of single-linkage clustering groups for every threshold value (*t*) from 1 to 1,628 allelic mismatches (Figure S5). A first optimal plateau of the silhouette coefficient (S*t*) was identified at 194/1,628 (11.9%), which partitioned the population in 42 clusters, and which was used to define clonal groups (CG). A second S*t* optimum at 940/1,628 (57.7%) was chosen based on the combination of the S*t* value and the corresponding distribution of allelic mismatches, which partitioned the isolates into 17 different groups, defined here as sublineages (SL). For continuity of the nomenclature, both CG and SL single-linkage classification groups were named by inheriting the number from the most common 7-gene sequence type (ST) in that group (Figure S6); when more than one CG uniquely or mostly comprised the same ST, incremental numbers starting at 10,000 were attributed to the smaller groups (e.g., for two groups predominantly made of isolates with ST326, CG316 and CG10000 were attributed).

### Host and geographic distribution of *C. ulcerans* sublineages and clonal groups

Three highly prevalent sublineages were observed: SL325 (n=211, 36.3%), SL331 (n=205, 35.2%), and SL339 (n=64, 11%). SL331 comprised mainly ST326, ST328, ST331, ST332 and ST358 isolates, SL325 mainly ST325, ST337 and ST543, whereas SL339 was mostly made up of ST339 isolates (Figure S6). The majority of the isolates belonging to these sublineages are *tox*-positive (80%), with the most common *tox* alleles being tox_33, tox_34, tox_49, and tox_50 (see below; note that the *tox* allele numbering includes alleles from *C. diphtheriae* too); these alleles are considered as functional, as tested isolates were Elek-positive, and the predicted translated protein does not contain frameshift mutations or insertion/deletion events leading to early stop codons. Two among the three major *C. ulcerans* sublineages were detected in more than one continent: SL325 in Europe (95.7%), Asia (3.8%), and Africa (0.5%), and SL339 in Europe (93.8%), Africa (4.7%), and South America (1.6%). All other sublineages were detected uniquely in one continent (Europe, n=13; South America, n=2), which could be an effect of their lower sampling sizes rather than a lack of large geographic distribution.

Clonal groups are finer-scale subdivisions of the population than sublineages. Whereas some sublineages comprise only one (major) clonal group (e.g., SL339), others are more heterogeneous (e.g., SL325 and SL331), and their clonal groups differ importantly in their genomic and biological characteristics.

Four clonal groups were highly represented: CG331, CG583, CG339 and CG325. CG325 (n=109) corresponded to an outbreak cluster among a group of dogs (2021-2022), which included isolates of ST325 and one of its single-locus variants, ST805. All isolates in this clonal group were *tox*-positive (allele tox_49), and the toxin gene corresponded to a single-gene contig (“orphan” toxin, see below). In contrast, CG543 (also belonging to SL325) were either *tox*-negative or *tox*-positive, in which case the *tox* gene was carried by a prophage (Figure 1). In CG339, another major clonal group (n=64), none of the genomes carried the *tox* gene. This clonal group was first reported in 2015 and was sampled in Europe, South America and Africa, and it was mostly isolates from animals (Figure 1).

Clonal group CG331 (n=96) comprised only *tox*-positive isolates, with only one exception; this *tox*-negative genome probably corresponds to a secondary loss (Figure 2A). The isolates from this clonal group were isolated both from humans and from animals (Figure 1). Interestingly, this clonal group, first reported in 2008 (Figure 2C) and which is geographically limited to Europe (France, Belgium, Germany and the UK), is associated with several toxin alleles (tox_34, n=69; tox_51, n=19; tox_64, n=1; and tox_3, n=1; for five isolates, it was not possible to determine the toxin allele, as the gene was fragmented, straddling multiple contigs) despite being genetically highly homogeneous. In this clonal group, the toxin was always carried by the *C. ulcerans* pathogenicity island (PAI), except for the toxin allele 3, which was carried by a prophage.

**Figure 2.**
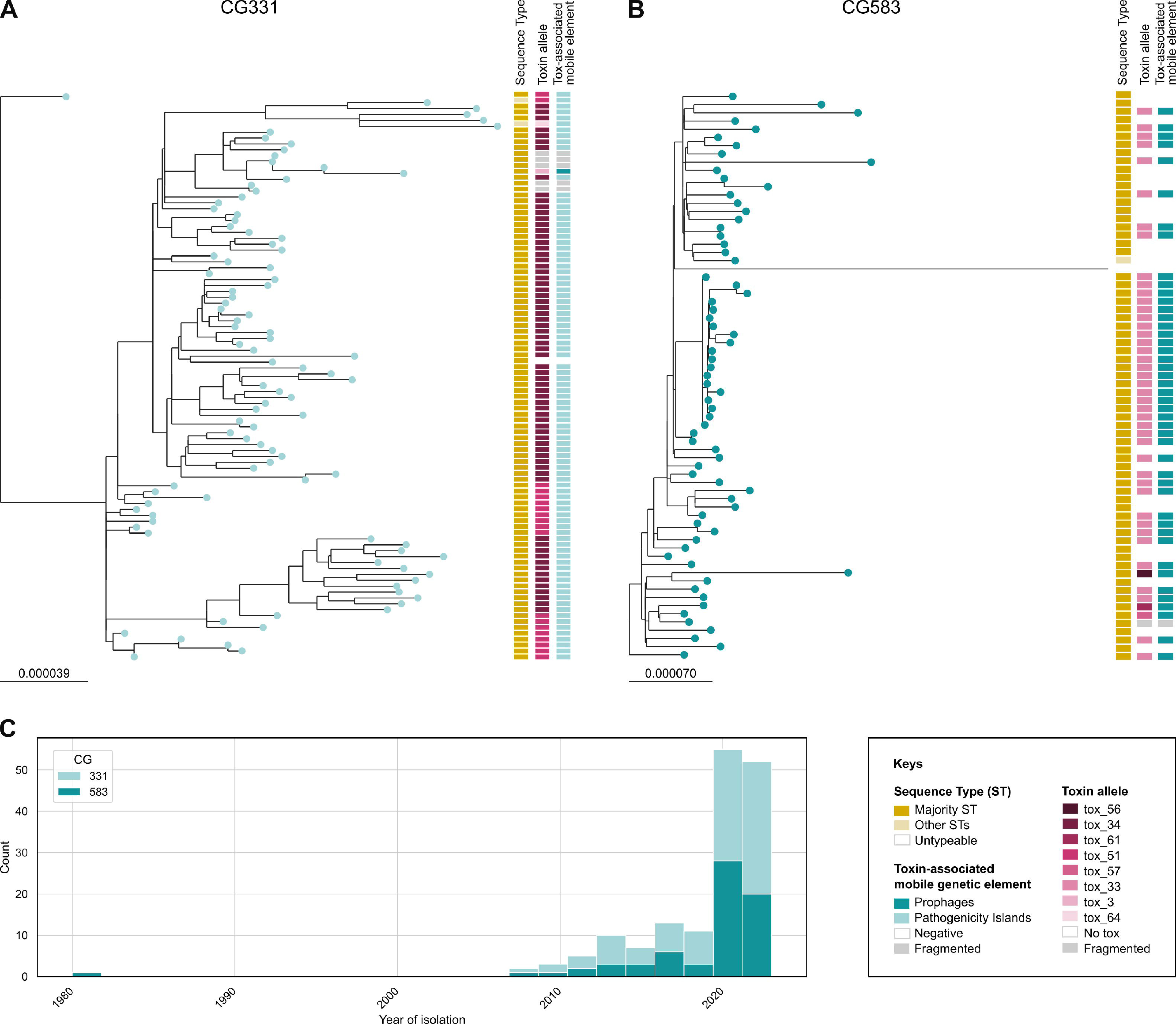
Phylogenetic trees of two major clonal groups (CG) of *Corynebacterium ulcerans*, (A) CG331; (B) CG583. Trees were rooted on the complete *C. ulcerans* phylogeny (Figure 1). The first external strip shows the majority ST in dark yellow (ST331 for CG331, and ST325 for CG583) and other STs (ST889 and ST933 for CG331, and ST583 and ST782 for CG583) in lighter yellow. The second and third external strips show the toxin alleles and the toxin-associated mobile genetic elements, respectively; (C) Timeline of isolation of CG331 and CG583 isolates.

CG583 (n=67) (Figure *2*B) is the second most prominent clonal group within SL325 (after CG325) and mainly comprises ST325 isolates, which were common both in humans and animals (Figure 1). In our dataset, this clonal group is restricted to European countries (i.e. France, Belgium, and Germany) and one isolate dates back to 1980 (Figure *2*C), while all other isolates were collected from 2008, as for CG331. In contrast to this and to other major clonal groups in *C. ulcerans*, which were either fully *tox*-positive or fully *tox*-negative (Figure *1*), isolates of CG583 were variably *tox*-positive (68.7%, n=46) or negative (31.3%, n=21).Figure *2* In this clonal group, the toxin was uniquely carried by a prophage. As *tox-*negative isolates were interspersed with *tox*-positive ones along the phylogeny, several loss events from *tox*-positive ancestors (Figure *2*B), or several acquisitions of *tox* by horizontal gene transfer (HGT) into previously tox-negative strains, must have occurred. The phylogenetic pattern of *tox* presence indicates that this prophage might be able to excise or invade frequently. A similar situation can be observed in CG543, a smaller clonal group of SL325 (Figure *1*).

### cgMLST variation within case clusters *versus* sporadic isolates

To calibrate cgMLST variation with respect to epidemiological events, the variation of cgMLST alleles was evaluated within groups of isolates reported as being epidemiologically related (i.e., a described cluster of cases, here “case clusters”) and compared to variation among sporadic isolates (i.e., isolates considered as being epidemiologically unrelated based on available source information). Twenty case clusters were identified (Table *1*), six of which were from the French NRC data (n=6), and 14 of which were from publicly available genome sequences and related metadata from different countries (Belgium, Spain, Germany, and the UK) (Figure *3*B). For four clusters reported in Belgium, only one genome sequence per cluster was available, and consequently, these clusters could not be analysed. The analysed case clusters mostly involved humans and their contact animals, mostly pets (dogs and cats), and they comprised isolates from various STs and carrying different toxin variants (Table *1*). We found a range of 0 to 19 cgMLST allelic mismatches (i.e., allele differences) among case clusters, and a mean variation of >1100 allele differences (Figure *3*A).

**Figure 3.**
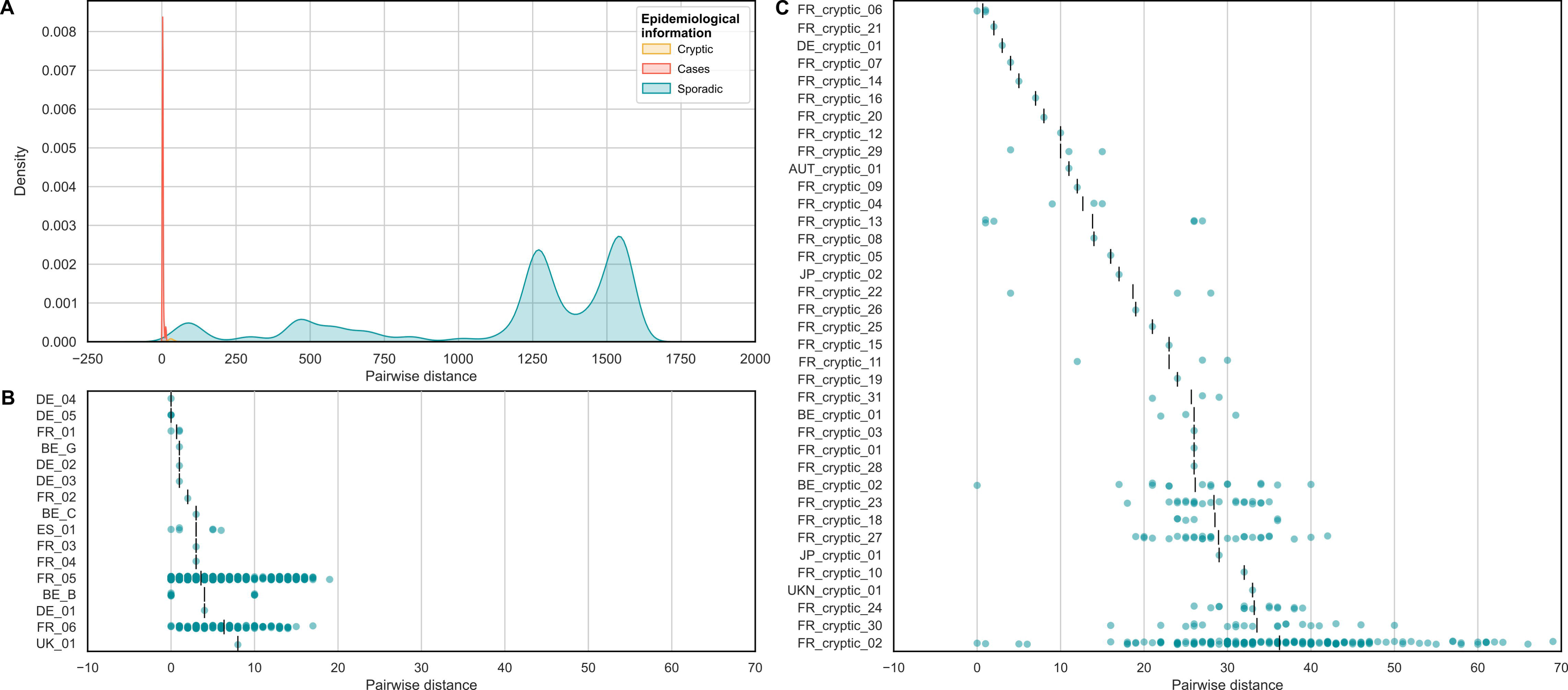
(A) Density plot showing the pairwise distances between cgMLST alleles among sporadic isolates, cryptic clusters and case clusters; (B) Pairwise distances among genomes within each case cluster, excluding those with only one genome available (Table 1); (C) Pairwise distances among genomes within each cryptic cluster, defined xas single linkage groups with ≤25 cgMLST mismatches. In B and C, each dot represents the pairwise distance between two isolates, whereas the vertical black line indicates the mean for each cluster.

### cgMLST genetic clusters reveal cryptic transmission of *C. ulcerans*

A threshold of ≤25 cgMLST mismatches was used to classify the 582 *C. ulcerans* genomes into single linkage groups, which we named genetic clusters. This number of mismatches was chosen based on the distribution of cgMLST allelic mismatches in the overall dataset (Figure *3*A) and the variation observed within clusters (Figure 3B). We then defined cryptic clusters as genetic clusters with no prior or subsequent evidence of epidemiological links. Case clusters (Figure *3*B) show a lower genetic heterogeneity compared to cryptic clusters (Figure *3*C), with cgMLST allelic mismatches always below 20 for the former, but extending to just under 70 in the latter (note that the single linkage definition of genetic clusters implies the possibility to two isolates having more than 25 mismatches, as long as they are linked by intermediate isolates with ≤25 mismatches). However, the majority (n=16) of cryptic clusters had a genetic heterogeneity comparable to that of case clusters (<20 mismatches among all isolates in the cluster). This could correspond to true case clusters for which there is simply a lack of evidence for epidemiological links, suggesting the potential utility of cgMLST for identifying possible undetected transmission.

We detected 38 cryptic clusters (Figure 3C; Table S1), all of which were single-country clusters. Nineteen of these were single-host species clusters (n=11 human-only, n=8 animal-only), whereas eighteen involved multiple host species. Among the latter, nine included at least one human and one domestic animal isolate (dogs, cats or horse), whereas ten included uniquely animal isolates from different species (both domestic and wildlife).

Interestingly, three cryptic clusters were linked to known case clusters. In two cases, they involved isolates that were collected in the same geographical areas as the epidemiologically related isolates, with a maximum distance of 60 km between isolates and a time range of 6 years for the first cluster, and of 50 km and 9 years for the second cluster (Figure S7). The third cryptic cluster included an isolate from a homeless patient that was linked to a known case cluster of two domestic rats from a nearby region that had been purchased from the same pet store (Museux et al., 2023) (Figure S7).

Most cryptic clusters originated from France (n=31), and so do those with the largest number of isolates, which is a direct consequence of the higher availability of sequence data from France. Interestingly, three among the major cryptic clusters from France (FR_cryptic_02, n=19 isolates; FR_cryptic_23, n=7 isolates; and FR_cryptic_27, n=8) showed restricted geographic distributions. The first was uniquely reported in the regions of the North of France, the second in the North-East, and the third was mostly found in the South (Figure S8). These data illustrate the power of cgMLST to uncover cryptic transmission and local geographic spread of *C. ulcerans* strains.

### Diphtheria toxin gene variation in the *C. diphtheriae* Species Complex and within *C. ulcerans*

The diphtheria toxin is the most important virulence factor of *C. ulcerans* and *C. diphtheriae*, and it is highly similar in these two species. A total of 399 *C. ulcerans* genome assemblies (68.6%) carried a complete *tox* gene. For thirteen additional *tox-*positive genomes, diphtOscan reported only partial gene hits, which corresponded to toxin gene sequences straddling two contigs. Twenty new toxin alleles were detected from *C. ulcerans* genomes and added to the *tox* gene sequence allele definitions in the BIGSdb database (allele numbers tox_46 to tox_65).

We compared the diversity of the diphtheria toxin gene from *C. ulcerans* with tox alleles from the entire CdSC, including sequences from *C. diphtheriae, C. ramonii, C. silvaticum*, and *C. pseudotuberculosis* (n=63 alleles in total). Four sequence clusters, here named “toxin families”, were identified (Figure 4) and named them based on the main bacterial host species associated with alleles in each cluster: Diphtheriae Toxin Family (DTF), Ulcerans Toxin Family (UTF), Silvaticum Toxin Family (STF) and Pseudotuberculosis Toxin Family (PTF). Some toxin alleles are unique to one bacterial host species, as was the case for alleles within the STF and PTF families, and for most DTF alleles, whereas others were found in more than one species, for example, DTF tox_2 and tox_3, and eight UTF alleles (Figure 4).

**Figure 4.**
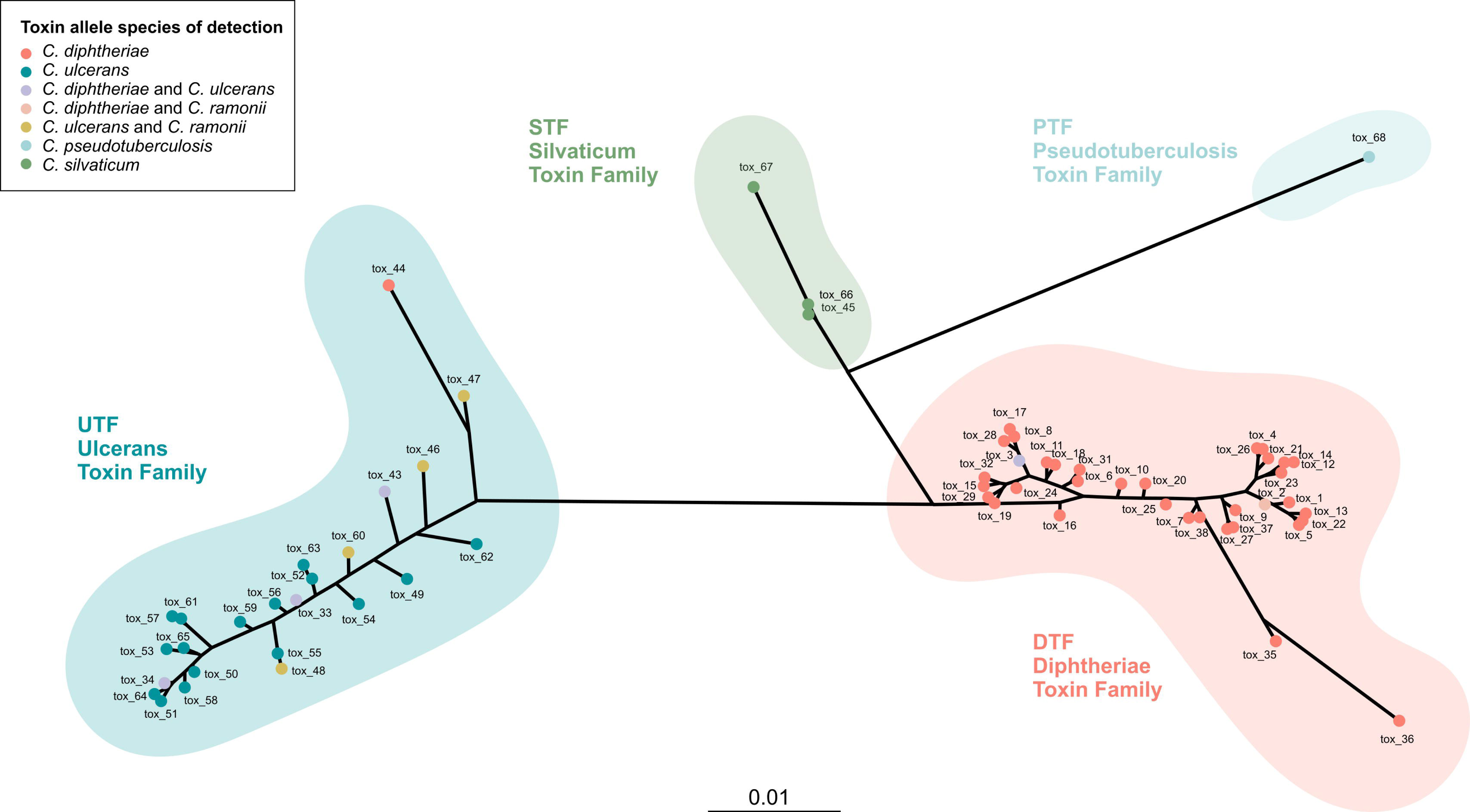
Unrooted phylogenetic tree of toxin alleles from all *tox*-carrying species of the *Corynebacterium diphtheriae* species complex (CdSC): *C. diphtheriae, C. ulcerans, C. ramonii, C. silvaticum,* and *C. pseudotuberculosis*. Toxin alleles clustered into four main “toxin families”, named after their main bacterial host species: Diphtheriae Toxin Family (DTF), Ulcerans Toxin Family (UTF), Silvaticum Toxin Family (STF) and Pseudotuberculosis Toxin Family (PTF). Some alleles are unique to one bacterial host species (e.g., STF, PTF and the majority of DTF alleles), whereas others are found in more than one species (e.g. DTF tox_2 and tox_3, and eight alleles of the UTF).

In *C. ulcerans*, the most commonly observed alleles were tox_33 (in n=57 genomes), tox_34 (n=78), tox_49 (n=108), and tox_50 (n=104). Toxin gene alleles were each associated with a single *C. ulcerans* sublineage, with three exceptions (tox_3 in SL325 and SL331; tox_43 in SL428 and SL329; tox_50 in SL331 and SL514) (Figure *5*A). Conversely, whereas certain sublineages were associated uniquely with one toxin allele (e.g., SL428 and tox_43, SL690 and tox_52), some SL and CG harboured more than one (e.g., SL325 n=9 *tox* alleles, SL331 n=8 *tox* alleles, CG331 n=4 *tox* alleles, CG337 n=3 *tox* alleles), suggesting several events of mobile genetic elements having imported these *tox* variants into these sublineages.

**Figure 5.**
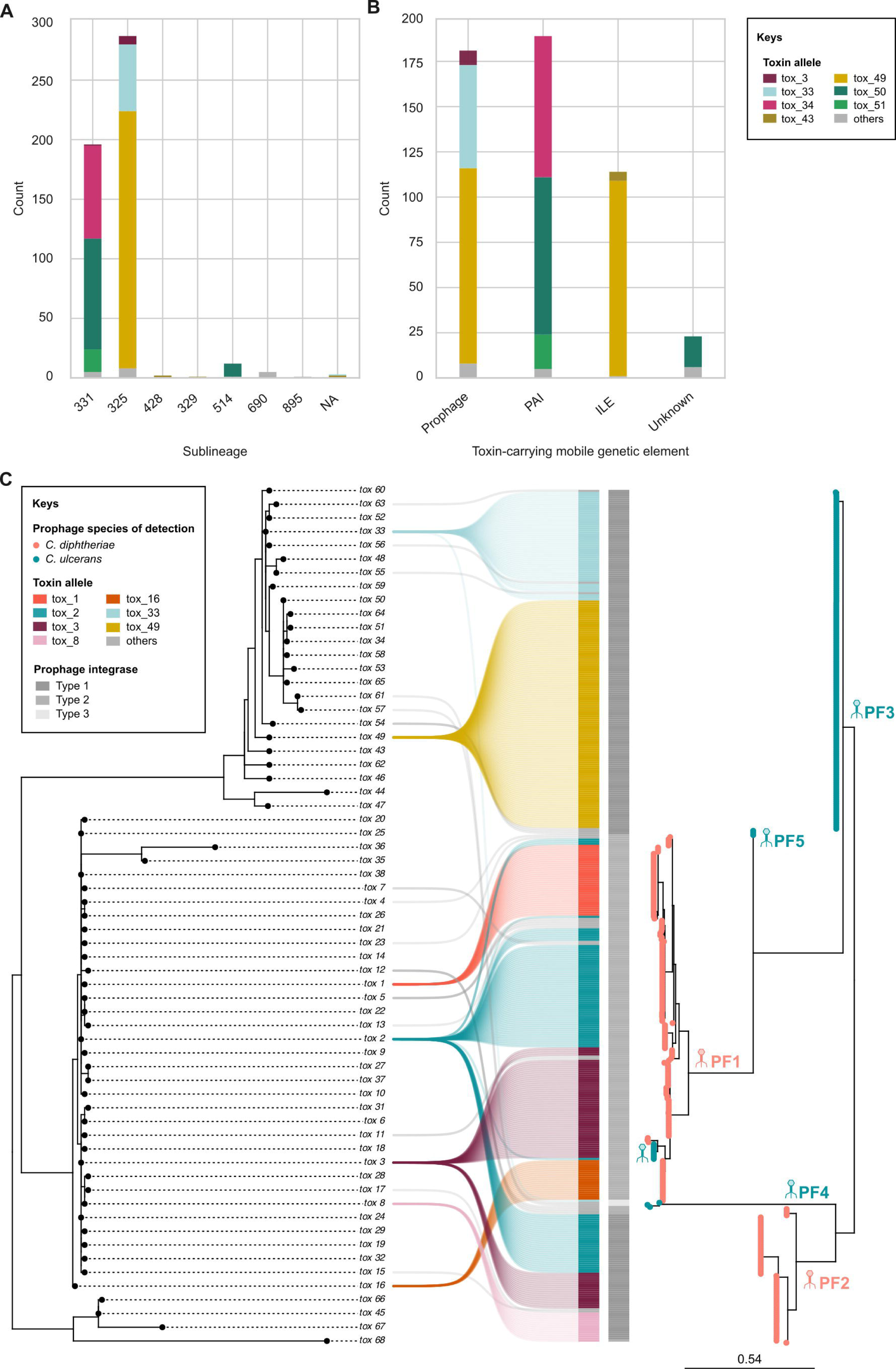
A) Bar plot showing the main toxin alleles found in *C. ulcerans* and their sublineages; NA: not assigned B) Bar plot showing the main toxin alleles found in *C. ulcerans* and the mobile genetic elements (MGE) carrying them; unknown MGE are due to high genome fragmentation; PAI: pathogenicity island; ILE: integron-like element. C) Phylogeny of *C. ulcerans* and *C. diphtheriae* toxin-carrying prophages (n=176 and n=232, respectively) and their toxin alleles, with main alleles showed in colours. Prophage families (PF) are indicated on the branches of the prophage phylogeny (PF1-PF5), in which leaves are coloured based on bacterial host species of detection.

### Mobile genetic elements associated with the diphtheria toxin gene

To define the genetic elements that carry the *tox* gene in *C. ulcerans*, we first sought to clarify the location of orphan toxins (i.e., contigs with only the *tox* gene), which were found in all isolates of CG325 (108/399, 27.1% of all *tox-*positive isolates), a clonal group that corresponds to an outbreak among dogs. We performed long-read sequencing of representative CG325 genomes. Strikingly, the resulting high-quality hybrid assemblies revealed the presence of two copies of the toxin gene (tox_49 allele for both), one carried by a 35,770 bp prophage, and the other by an integron-like element (ILE, of 3,230 bp). The ILE is inserted a few genes (∼8,000 bp) downstream of the prophage (Figure S9), and although IntegronFinder (Néron et al., 2022) did not identify it as such, the structure of this element, which is constituted solely by a tyrosine recombinase/integrase and the toxin, is similar to that of an integron. The presence of two identical copies of the *tox* gene explains the failure of Illumina data assembly, in which all *tox*-gene reads from both copies were collapsed into a single contig during assembly.

When considering the other *C. ulcerans* clonal groups and their *tox*-carrying elements, the diphtheria toxin was found to be approximately equally carried either by the previously-described PAI (189/399, 47.4% of the *tox-*positive genomes) or by prophages (181/399, 45.4%, including the CG325 prophages) (Figure *5*B). In six genomes (1.5% of the *tox-*positive genomes), the toxin was carried uniquely on an ILE. Together with the CG325 genomes, the *tox-*ILE was present in 28.6% of genomes. For thirty-six genomes, it was not possible to identify clearly which genetic element was carrying the toxin, due to genome assembly fragmentation.

The *tox*-carrying prophage sequences clustered into five main genetic groups, which we named here prophage families (PF1 to PF5; Figure *5*C). Four prophage families were associated uniquely with one bacterial host species (PF2 with *C. diphtheriae*, and PF3, PF4 and PF5 with *C. ulcerans*). In contrast, although prophages belonging to PF1 were primarily detected in *C. diphtheriae*, a PF1 sub-clade carrying allele tox_3 was observed in *C. ulcerans* isolates from SL325 and SL331, which is indicative of bacteriophage cross-species infection. We also observed that the three prophage families of *C. ulcerans* PF3, PF4, PF5 were associated uniquely with one sublineage: SL325, with the unique exception of one PF3 phage also observed in *C. ulcerans* SL514. In contrast, PF1 and PF2 were associated with several sublineages within *C. diphtheriae* and *C. ulcerans,* respectively (Figure S10), showing their ability to infect a broader range of *C. ulcerans* strains.

Vertical transmission of *tox*-carrying elements in *C. ulcerans* was evident from their distribution in the *C. ulcerans* phylogeny (Figure *1*), where *tox*-prophages and the PAI are each associated with one branch, mainly corresponding to SL325 (prophages) and SL331 (PAI). In addition, the PAI sequence is very conserved across the entire SL331.

Prophages and ILE showed different integrase amino-acid sequences, whereas all the PAI were characterized by the same integrase gene (Figure S11). Phage integrases 1 and 2 showed a high sequence similarity (90%), whereas phage integrase 3 was quite distinct from the others, with only ∼33% of amino-acid sequence identity. Regarding ILE integrases, integrases ILE-1 and ILE-2 were identical except for their sequence length (150 AA and 104 AA, respectively), whereas integrase ILE-3 shows a lower sequence identity (82%). Prophages with integrase 1 and the PAI integrase shared the same insertion site (Figure S11), which likely explains why these elements appear as mutually exclusive in their respective phylogenetic branches. ILE-1 and ILE-2 integrases and the PAI integrase were more similar to phage integrases 1 and 2, than was phage integrase 3.

### Other virulence genes

The phospholipase D (PLD, a sphingomyelinase) is another important virulence factor of *C. ulcerans*. We observed that 557 genomes (95.7%) carried the PLD exotoxin gene *pld*, with only SL349 being negative for this gene.

The *rbp* (ribosomal-binding protein) gene, another virulence factor described in the CdSC, was carried uniquely by one genome (809, an ST338 which is the reference genome for *C. ulcerans*).

## Discussion

To provide a reproducible, standardized, and high-resolution genotyping method for *C. ulcerans* strains, we developed a cgMLST scheme specific to this emerging pathogenic species. By applying this scheme to a large dataset of available and newly generated genome sequences, we illustrate how this novel tool will facilitate the study of *C. ulcerans* diversity and evolution, and how it will enable tracing its spread and transmission patterns across countries. Taking into account population structure, we defined groups of genetically similar genomes at different phylogenetic depths: from the deep subdivisions of sublineages (SL) (cgMLST mismatch threshold of 940), to the progressively shallower ones of clonal groups (CG) (194 mismatches) and genetic clusters (25 mismatches). To ensure readability of the new SL and CG nomenclature, we attributed identifiers inherited from the existing seven-gene MLST nomenclature using a previously described inheritance algorithm (Hennart et al., 2022). The cgMLST scheme, as well as the SL and CG nomenclatures, are publicly available on the diphtheria database in BIGSdb-Pasteur.

Availability of *C. ulcerans* genomic data from outside Europe is scarce, and the picture provided by this work is influenced by a bias towards French isolates. Future genomic studies of *C. ulcerans* from other countries will likely enhance its overall reported genetic diversity. Despite this, CG that comprise isolates from different countries (e.g., CG337, CG328, CG514), some of which far from Europe (e.g., Japan), do not show deep branching separating isolates from distinct geographical origins; hence, major clonal groups found in France might be also represented in other countries. Excluding case clusters, the *tox-* negative CG339 and the two *tox-*positive CG331 and CG583 (Figure *2*) appear as the dominant groups in the population, two of which (CG583 and CG339) were detected in multiple continents. All frequent CG are associated with three or more host species, including humans, domestic and wild animals; this provides population genomics insights into the zoonotic character of *C. ulcerans* sublineages. The population structure of *C. ulcerans* is strikingly different from that of *C. diphtheriae*, whose population structure shows deep branching of multiple (hundreds) of sublineages, and where no sublineage appears as being highly predominant (Guglielmini et al., 2018; Hennart et al., 2020). This points to a larger genetic pool of *C. diphtheriae* compared to *C. ulcerans*, even though the former is largely restricted to humans. Sampling of wild animals is expected to reveal currently hidden diversity of *C. ulcerans*, perhaps including sublineages that are not pathogenic for humans.

To date, the transmission dynamics of *C. ulcerans* have been poorly documented. Genomic surveillance can indirectly inform on transmission, by inference of the geographic distribution of single genotypes. We applied a 25 allelic mismatch threshold of cgMLST loci to define genetic clusters, which successfully included all reported epidemiological clusters of infections (case clusters). In addition to these, single-linkage clustering allowed us to detect several hidden clusters of infections, which show a low genetic heterogeneity comparable to that observed within the case clusters (Figure *3*B and Figure *3*C). This is different from what was observed previously in *C. diphtheriae*, in which most cryptic clusters show a higher heterogeneity compared to the case clusters (Guglielmini et al., 2018). Interestingly, when excluding single-host cryptic clusters, the remaining cryptic clusters include either human isolates and pets (dogs, cats, and horses), or pets and wildlife, with no cluster including both human and wild animal isolates. In addition, some cryptic clusters show a strong association with certain geographical areas, with dissemination over time across neighbouring regions (e.g., Northern France). Taken together, this data suggests that asymptomatic and symptomatic carriage of *C. ulcerans* in wild animal populations probably play an important role in its spread across vast geographical areas (Figure S8), with pets likely being at the centre of the transmission chain between wildlife and humans. However, when looking at the links between isolates within the current cryptic clusters of transmission, as well as at their years of isolation and geographical origin, it is clear how a lot of the diversity of the *C. ulcerans* population and of its transmission dynamics are currently not being captured. This is probably due to under-sampling, under-reporting and under-sequencing of isolates from the wildlife population, which currently impairs our understanding of the transmission and phylodynamics of *C. ulcerans*.

With this study, we provide fresh insights into the diversity of the *tox* gene and of its associated mobile genetic elements in *C. ulcerans*. Our phylogenetic analysis reveals how isolates of the two major sublineages, SL331 and SL325, are for the most part positive for the diphtheria toxin gene. Strikingly, the toxin was carried primarily by the PAI in SL331 and by prophages in SL325, suggesting an ancestral acquisition of these elements in the two branches. The PAI is well conserved across all clonal groups of SL331 and, differently from the prophage, which is lost multiple times within SL325, it does not seem to be easily excised. For the most part, toxin alleles were each associated with a single sublineage, with some exceptions, and this is true also for their associated MGE (see our Microreact project interactive exploration and details). Additionally, we report for the first time the presence of a small genetic element carrying the *tox* gene alongside an integrase (“integron-like” element, ILE), and the observation of two copies of the *tox* gene in *C. ulcerans*, in genomes from CG325, one carried by a full prophage, and the other by an ILE. Carriage of two toxin genes has so far only been observed in *C. diphtheriae*, in the toxin hyperproducer vaccine strain PW8, in which, however, two copies of the toxin are carried by two full prophages, a situation likely due to *in-vitro* evolution upon strain selection for toxoid vaccine production.

We detected twenty new toxin alleles in *C. ulcerans* and classified all toxin alleles into four "toxin families" (*C. diphtheriae* DTF, *C. ulcerans* UTF, *C. silvaticum* STF, and *C. pseudotuberculosis* PTF). Similar to our findings, one previous study on phylogenetic analysis of twelve *C. diphtheriae* and *C. ulcerans* isolates reported a division in two species-specific clades of their diphtheria toxin sequences (Otsuji et al., 2019), corresponding to the two groups now named DTF and UTF; however, in contrast to the previous study, here we show how some toxin alleles are not unique to one bacterial species, and how e.g., a toxin allele typical of *C. diphtheriae* (e.g., DTF tox_2 and tox_3) can be found in other species (e.g., *C. ramonii* and *C. ulcerans*, respectively) (Figure *4*), likely as a result of horizontal gene transfer via phage lysogeny. We also describe for the first time the existence of five *tox*-carrying prophage families in the CdSC, some unique to one bacterial species and others present in multiple species, similar to the toxin gene, pointing to cross-species infection and to cross-species toxin allele transfer. Transmission of the toxin in *C. ulcerans* appears to be both vertical (via PAI and phages) and horizontal (especially phage-mediated). These findings open interesting questions about the co-evolution of the toxin and its carrying genetic elements, also considering the introduction of the *C. diphtheriae* vaccine in the first half of the 20^th^ century. Despite several differences at the amino-acid level of the diphtheria toxin (DT) from *C. diphtheriae* and *C. ulcerans* in the receptor-binding domain and in the translocation region (Sing et al., 2003), the diphtheria toxoid vaccine has been shown to be also active against the diphtheria toxin produced by *C. ulcerans* (Möller et al., 2019). However, the impact of the acquisition of a toxin of the DTF via phage infection on the pathogenicity of *C. ulcerans* is still unknown and should be explored, especially considering the current epidemiological situation with rising *C. ulcerans* cases worldwide.

In conclusion, dominant clonal groups of *C. ulcerans* and their cryptic clusters, including those with a vast geographic dissemination, suggest potential transmission chains, with pets serving as intermediaries between wildlife and humans. Carriage and transmission between wild animals are poorly understood and under sampling of wild animal reservoirs currently hampers a comprehensive understanding of *C. ulcerans* transmission dynamics. The *C. ulcerans*-specific cgMLST scheme: i) adds to our genomic epidemiology toolkit to address the re-emergence of zoonotic diphtheria-like infections; ii) will facilitate harmonization of knowledge on *C. ulcerans*; iii) provides a common framework to study its diversity and transmission patterns at a global scale; and iv) it will facilitate the generation of new knowledge on its ecology, evolution and transmission. While many questions, such as the dynamics of toxin-MGE evolution, remain open, this study significantly contributes to our understanding of *C. ulcerans* epidemiology, genetic diversity and evolution.

## Data availability

All genome sequences published in this study were made available at BIGSdb-Pasteur (project id 45 at https://bigsdb.pasteur.fr/cgi-bin/bigsdb/bigsdb.pl?db=pubmlst_diphtheria_isolates). Supplementary material includes Table S1 (isolate metadata for strains used in this study) and a Supplementary appendix. A project with the phylogenetic tree, metadata and results file is available within Microreact at this link: https://microreact.org/project/pJf35xQkcs2HUznD67W7Be-corynebacterium-ulcerans-genomic-epidemiology-2024.

## Authors contributions

Conceptualization: Sylvain Brisse, Chiara Crestani; Methodology, Data Curation, Validation and Visualization: Chiara Crestani; Software: Martin Rethoret-Pasty; Formal Analysis: Chiara Crestani, Alexis Criscuolo; Writing original draft: Chiara Crestani; Writing review & editing: Sylvain Brisse, Chiara Crestani; Supervision and Funding Acquisition: Sylvain Brisse.

## Acknowledgements

We thank the Mutualized Platform for Microbiology (P2M) for sequencing isolates from the French NRC using Illumina technology. This work used the computational and storage services provided by the IT Department at Institut Pasteur. The National Reference Center for Corynebacteria of the *diphtheriae* complex is supported financially by Institut Pasteur and Santé Publique France (Public Health France). This work was supported financially by the French Government’s Investissement d’Avenir grant Laboratoire d’Excellence Integrative Biology of Emerging Infectious Diseases (ANR-10-LABX-62-IBEID). We thank Dr Julie Toubiana for her clinical expertise and support to the French NRC of CdSC.

## Footnotes

1 The official number of *C. ulcerans* genomes available on the Genome page of NCBI in June 2022 is of 37; however, n=7 genomes were recently classified as belonging to a different species (*C. ramonii*) and therefore discarded at this stage, and NCTC13718 was discarded as, although it corresponded to a *C. ulcerans* genome, the NCTC number is attributed to a *Pseudomonas aeruginosa* isolate. Additionally, we included the public genome sequence of KL1017, from a case of bovine mastitis.

