## Supplementary appendix for "Genomic Epidemiology and Microevolution of the Zoonotic Pathogen *Corynebacterium ulcerans*"

### Figures

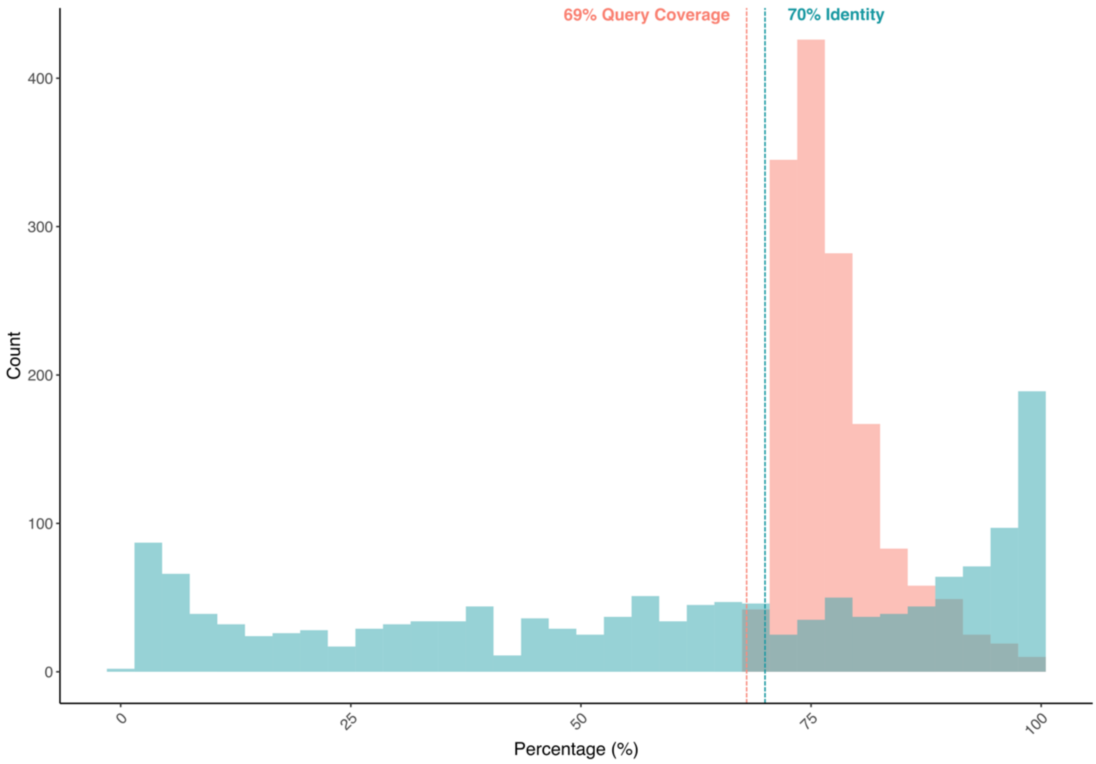


Figure S1 Distribution of percentage of identity (in blue) and of query coverage (in pink) for BLASTn pairwise comparisons between the *C. diphtheriae* cgMLST loci (n=1,305) and the filtered *C. ulcerans* cgMLST loci from chewBBACA (n=1,888).

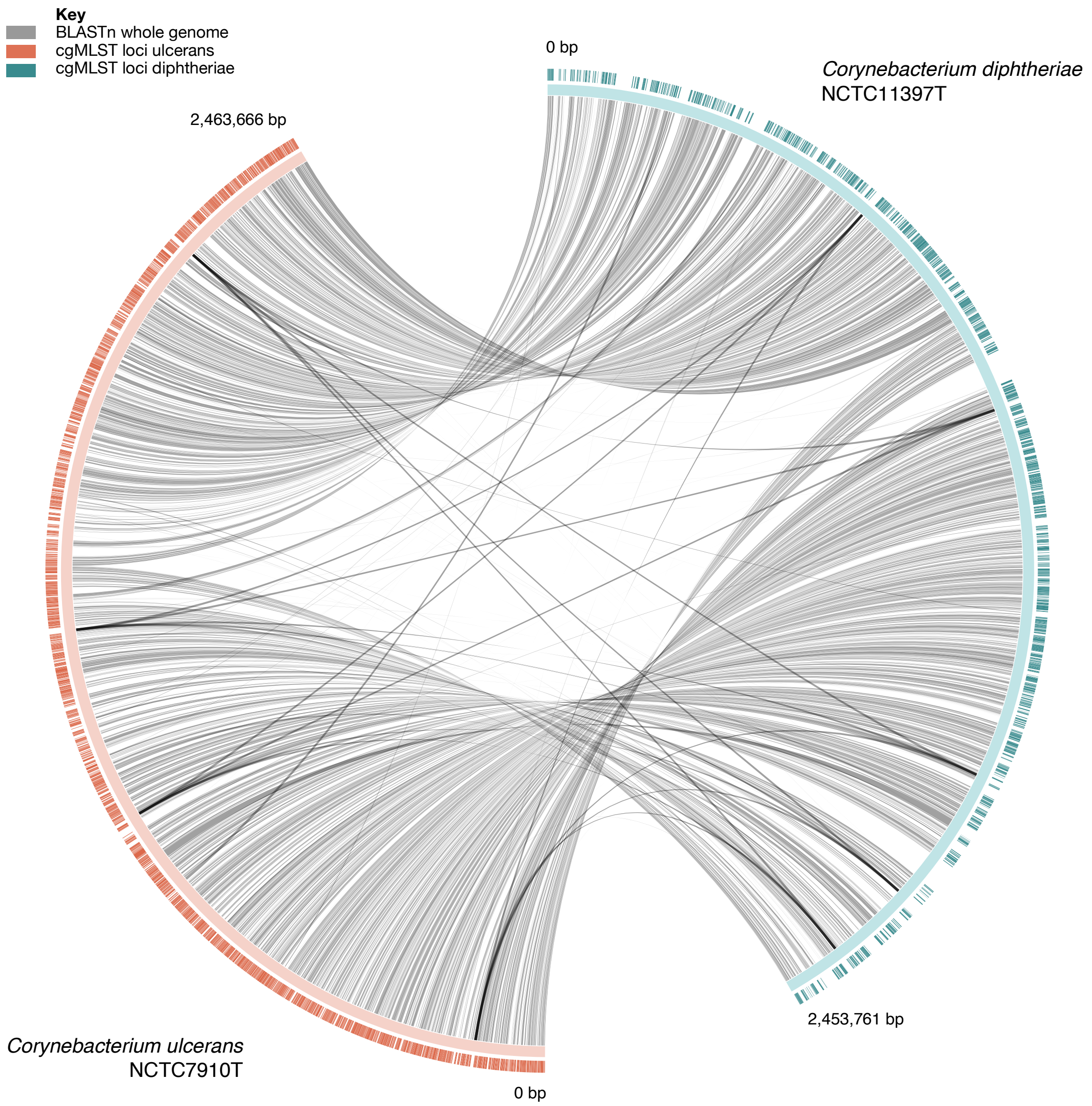


Figure S2 Circular plot representative of the two reference genomes for *C. ulcerans* (NCTC7910T) and *C. diphtheriae* (NCTC11397T) (continuous orange and green lines), of their genome identity (grey ribbons), and of the position of the genomes of their respective cgMLST loci. Alleles in common were found to be n=519 (BLAST ID >69%, query coverage >70%, 5% allele variation allowed).

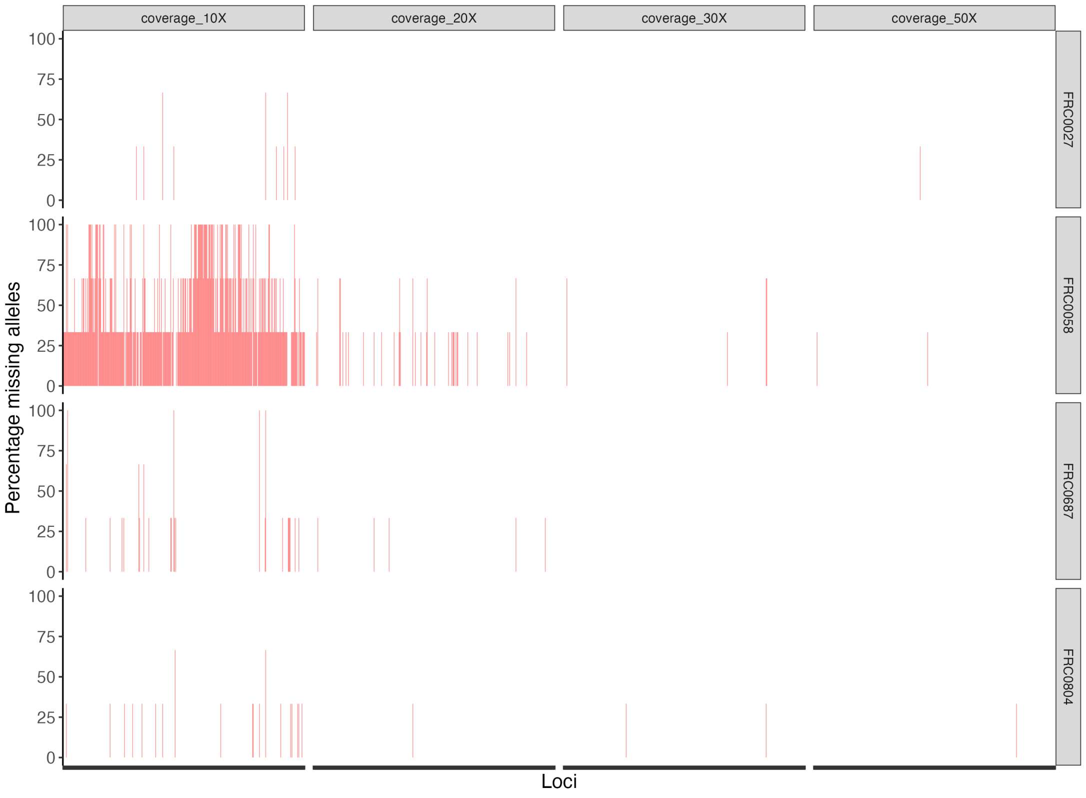


Figure S3 Allele call reproducibility at simulated low coverage depths for four *C. ulcerans* genomes (FRC0027, FRC0058, FRC0687, FRC0804). Assemblies built on simulated coverage (10X, 20X, 30X, 50X, 80X, 100X; in triplicate for each genome at each coverage) were tagged for existing alleles of the cgMLST scheme, to identify loci with non-reproducible allele calls at low assembly coverage depths. All alleles were called with no mismatches at 80X and 100X, whereas 9 unique loci showed non-reproducible allele calls at 30X and 50X. These were eliminated from the scheme, for a final set of 1,628 loci of the *C. ulcerans* cgMLST scheme.

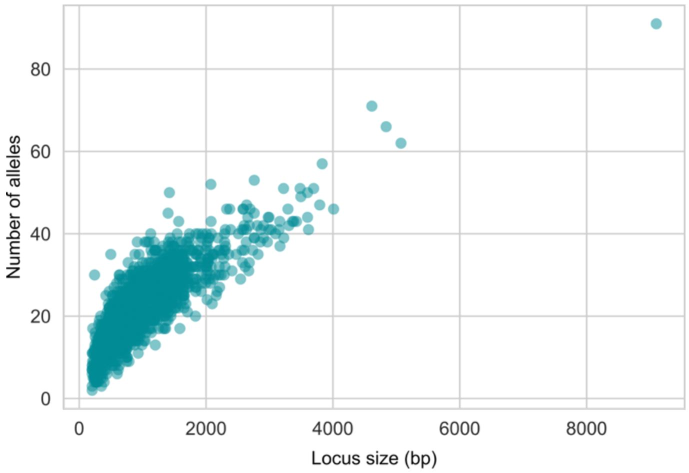

Figure S4 Scatterplot showing the relationship between the size of the loci in the cgMLST scheme of *C. ulcerans* and the number of alleles corresponding to that locus.

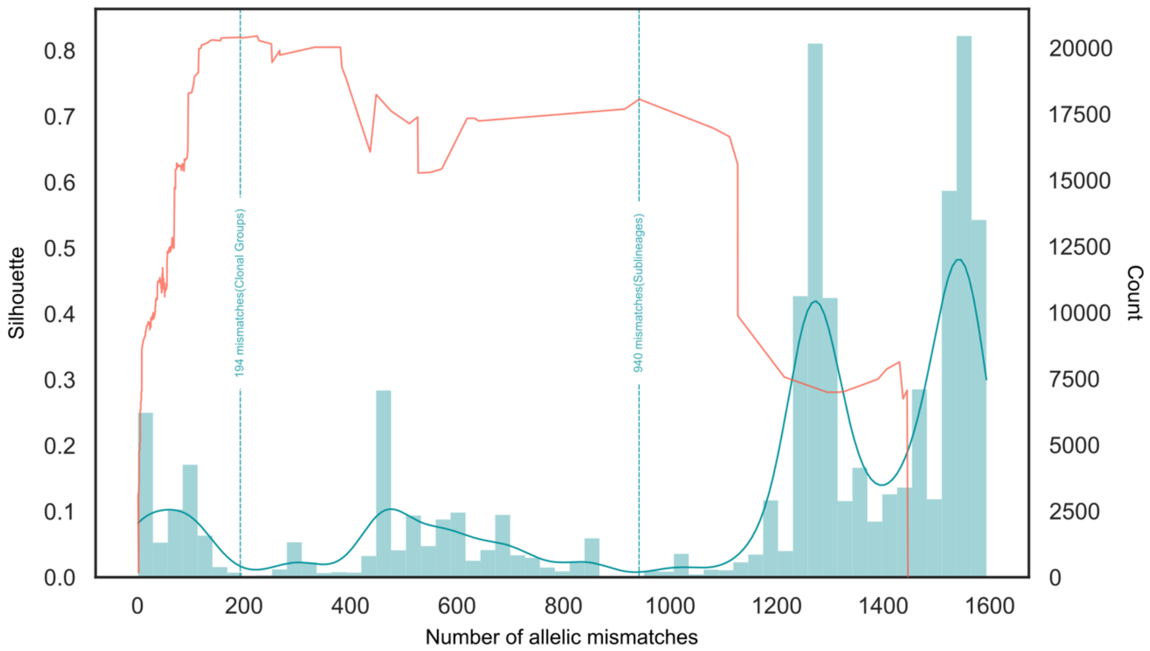


Figure S5 Frequency plot illustrating the number of cgMLST allelic mismatches between all pairs of *Corynebacterium ulcerans* genomes in the validation dataset, except for those with incomplete cgMLST profiles (n=5) and duplicated genomes (n=2). The silhouette coefficient ($S_t$) was plotted as a red line. The y-axis on the left shows values for $S_t$, while the one on the right shows the count for pairwise allelic mismatches. The two dotted lines correspond to the threshold for clonal group (194 mismatches) and sublineage (940 mismatches) definitions.

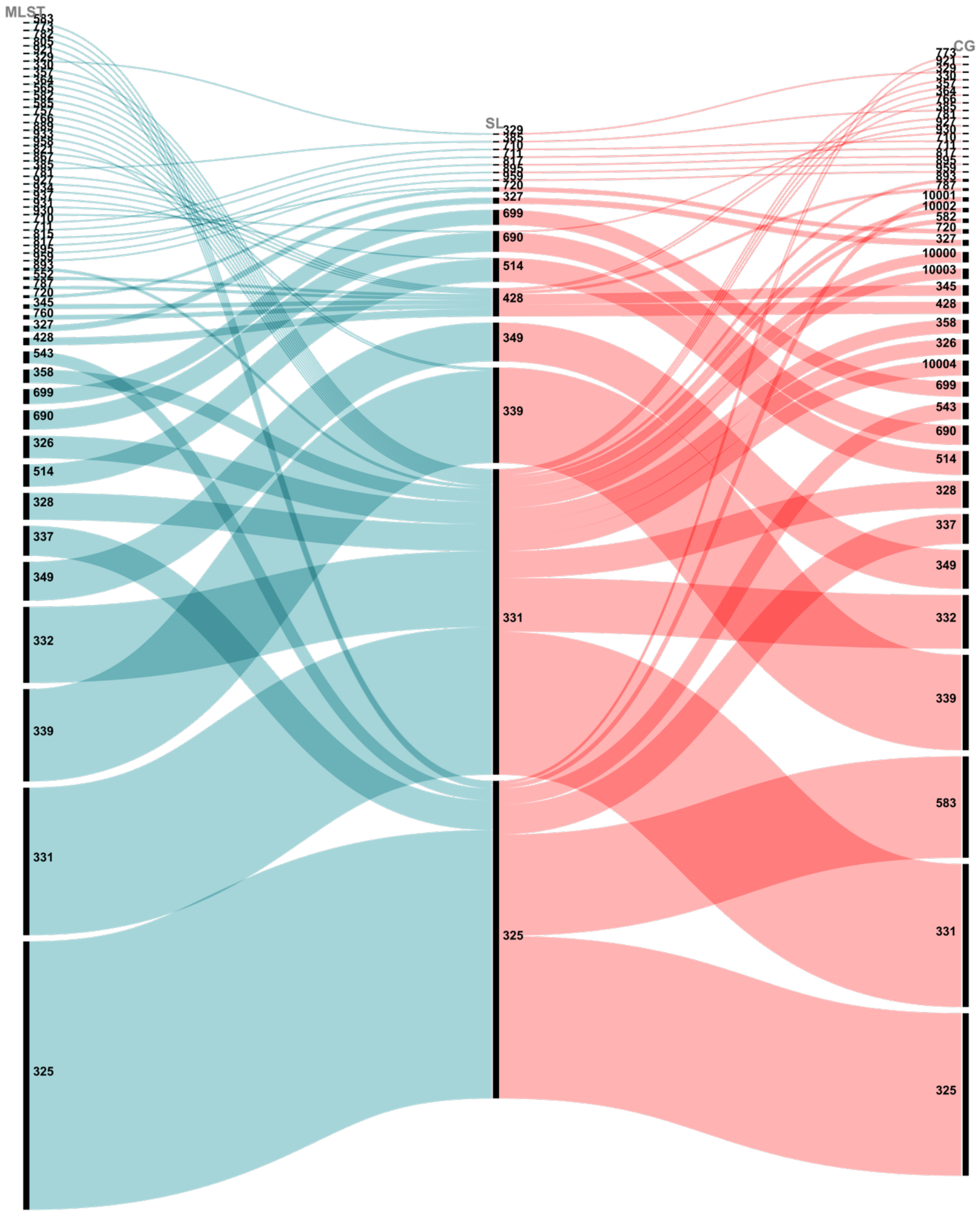


Figure S6 Alluvial diagram showing how *C. ulcerans* Sequence Types (ST – MLST) map to its sublineages (SL) and clonal groups (CG), and the inheritance of the 7-gene MLST nomenclature of SL and CG.

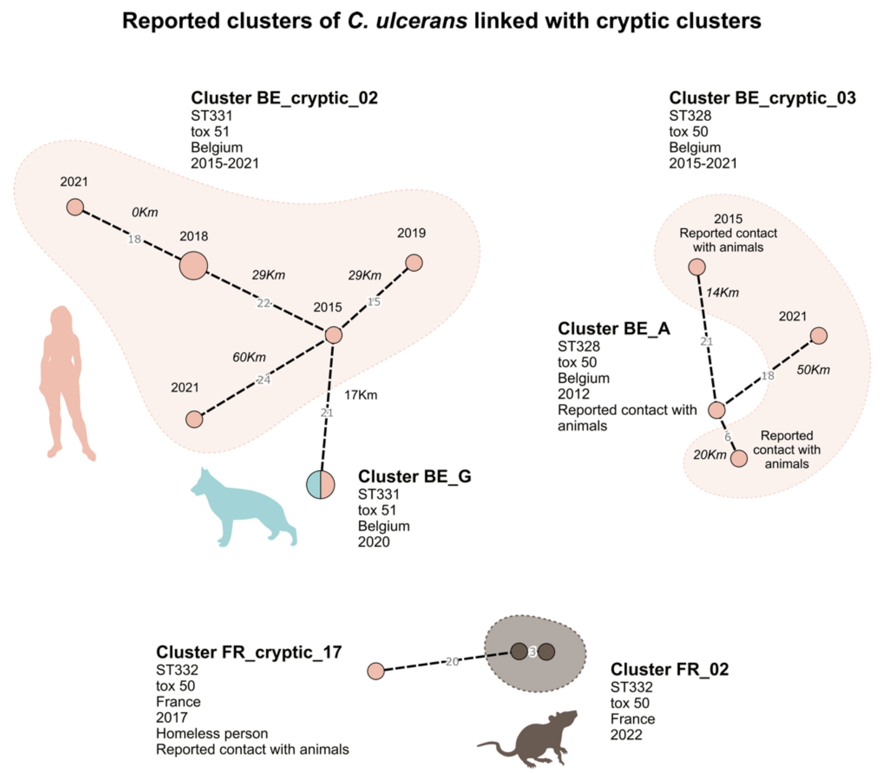


Figure S7 Three cryptic clusters linked to reported outbreak clusters detected in this work.

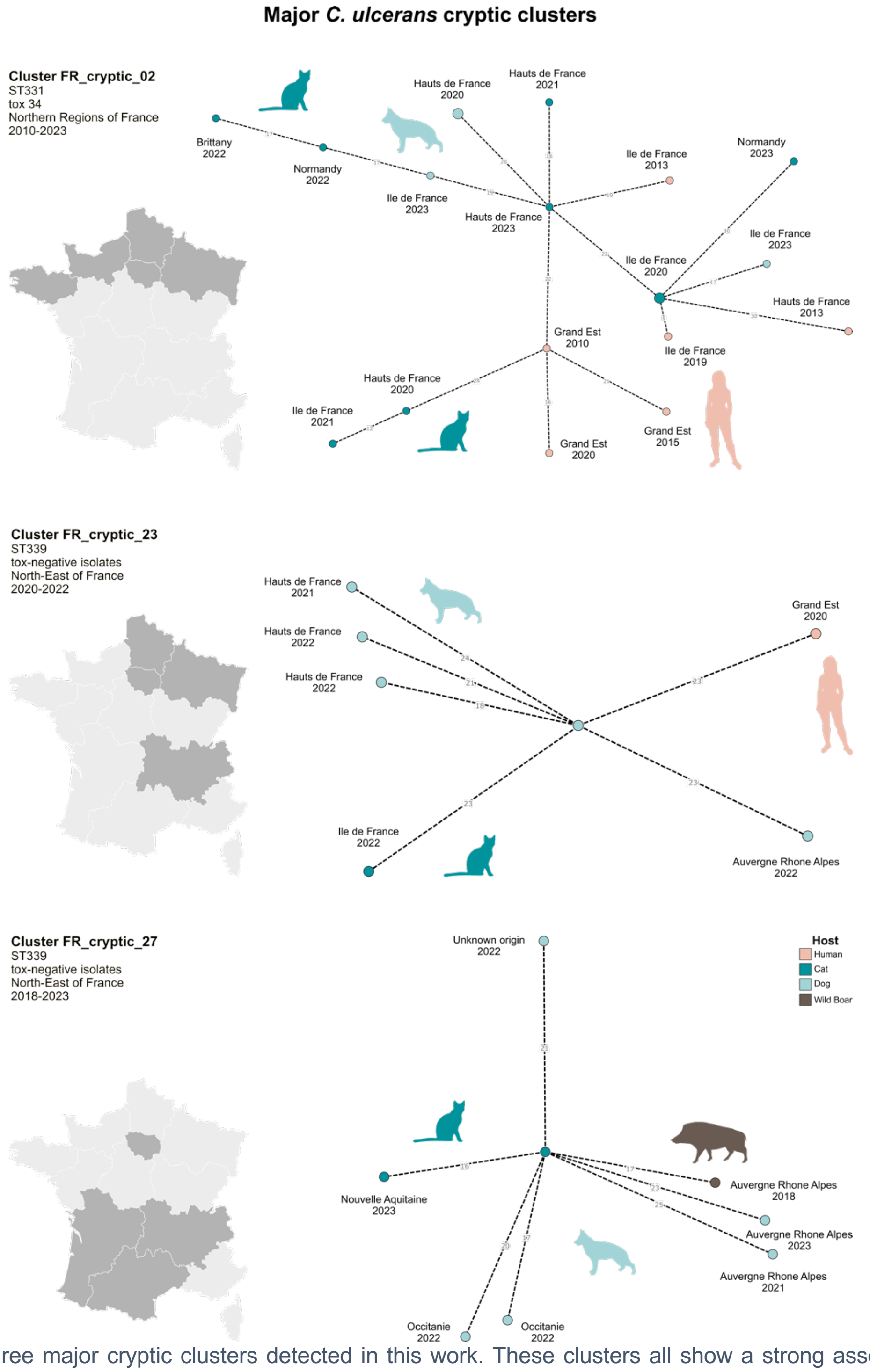


Figure S8 Three major cryptic clusters detected in this work. These clusters all show a strong association with distinct geographical areas.

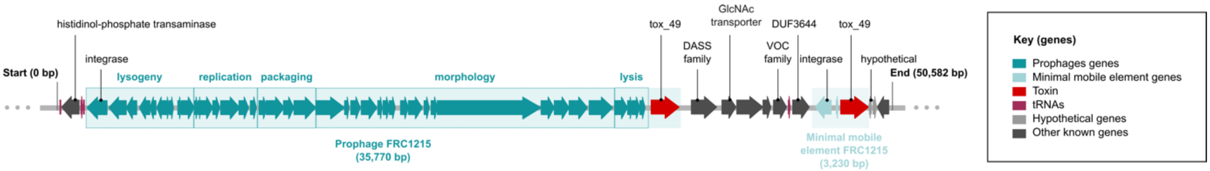


Figure S9 Map of a chromosomal region of a hybrid long and short-read assembly of isolate FRC1215, shown here as representative of CG325 genomes, all of which carried an orphan toxin in their short-read assembly. Long-reads helped in elucidating both the cause of the poor short-read assemblies (i.e. a double copy of the toxin), and the mobile genetic elements carrying the toxin (a prophage and a minimal mobile element).

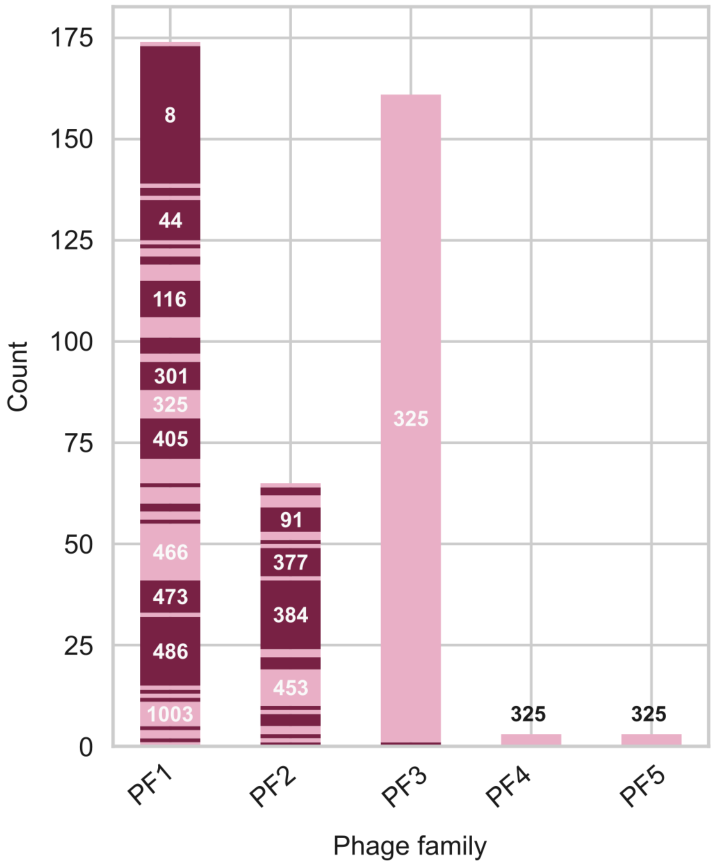


Figure S10 Bar chart showing the association of prophage families (PF) with sublineages (SL) of *C. diphtheriae* and *C. ulcerans*. PF3, PF4 and PF5, which were only detected in *C. ulcerans* genomes, are uniquely associated with one lineage (SL325), except for one phage from PF3, whereas PF1 and PF2 are associated with multiple phylogenetic lineages in *C. diphtheriae* and *C. ulcerans.*

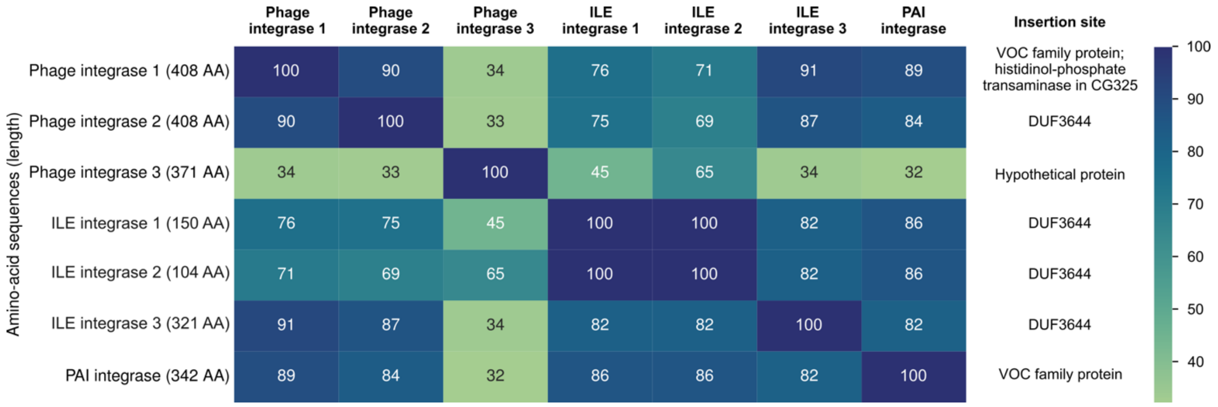


Figure S11 Heatmap of the amino-acid sequence identities of integrase genes belonging to *tox*-carrying mobile genetic elements (MGE) detected in this study. Insertion sites where each of these were detected are indicated in the last column; most integrases are site-specific (one insertion site only), except for phage integrase 1, which had a different insertion site in GC325 genomes (Figure S9). Some of these integrases share the same insertion site (e.g. phage integrase 2, with ILE integrase 1, 2 and 3; and phage integrase 1 with the PAI integrase), which likely explains why these elements appear mutually exclusive.
